# Computational Fingerprints: Modeling Interactions Between Brain Regions as Points in a Function Space

**DOI:** 10.1101/2021.09.28.462195

**Authors:** Craig Poskanzer, Stefano Anzellotti

## Abstract

In this paper we propose a novel technique to investigate the nonlinear interactions between brain regions that captures both the strength and the type of the functional relationship. Inspired by the field of functional analysis, we propose that the relationship between activity in two different brain areas can be viewed as a point in function space, identified by coordinates along an infinite set of basis functions. Using Hermite Polynomials as basis functions, we estimate from fMRI data a truncated set of coordinates that serve as a “computational fingerprint,” characterizing the interaction between two brain areas. We provide a proof of the convergence of the estimates in the limit, and we validate the method with simulations in which the ground truth is known, additionally showing that computational fingerprints detect statistical dependence also when correlations (“functional connectivity”) is near zero. We then use computational fingerprints to examine the neural interactions with a seed region of choice: the Fusiform Face Area (FFA). Using k-means clustering across each voxel’s computational fingerprint, we illustrate that the addition of the nonlinear basis functions allows for the discrimination of inter-regional interactions that are otherwise grouped together when only linear dependence is used. Finally, we show that regions in V5 and medial occipital and temporal lobes exhibit significant nonlinear interactions with the FFA.

## Introduction

The use of linear and nonlinear models for the analysis of neuroimaging data is at the center of a lively debate [1]. On one hand, proponents of linear models argue that nonlinear models can lead to overfitting issues given the amount of data that is typically available [2]. Linear models have been found empirically to yield insights about the brain when used for pattern classification [3], representational similarity analysis [4], and functional connectivity [5]. On the other hand, proponents of nonlinear models argue that linear models are not biologically plausible: firing rates of single neurons are integrated nonlinearly within dendrites [6, 7], and nonlinear transformations are essential to perform many of the tasks humans need to solve. Thus, while linear models might be effective to test whether a brain region encodes a given set of features, they might fall short of capturing interactions between brain regions with a complexity sufficient to enable the understanding of cognitively relevant computations.

More broadly, research on the statistical dependence between the responses in different regions (“functional connectivity”) has focused on studying *whether* given pairs of brain regions interact; however, there is a need for methods that can be used to investigate *how* they interact – to distinguish between different kinds of mappings that transform information from brain region to brain region. Even interactions between brain regions displaying similar strengths of functional connectivity could belie very different nonlinear computations.

A recurring criticism of nonlinear models is based on the difficulty to interpret them. In decoding analyses, linear models make it easier to distinguish the contribution of neural information processing up to the brain region whose responses are being measured from the contribution of the decoder applied to extract information from that brain region (see [8, 9]). By contrast, nonlinear decoders can transform the neural responses they receive as inputs to an extent that might lead to ambiguity about the nature of representations in the brain region that is being investigated. To illustrate this point with an example, if we could use any nonlinear decoder, and we had noiseless data from every single neuron in early visual cortex, we should be able to use these data to perform view-invariant object classification. After all, the brain itself can perform view-invariant object classification using early visual cortex responses as input. However, this finding would not support the conclusion that early visual cortex encodes view-invariant representations of objects, because the nonlinearities in the decoder would have likely been necessary for view-invariance to occur.

This criticism of nonlinear decoders is largely motivated by the “standard” analysis strategy used in the literature. This standard strategy consists of training one model to achieve the highest possible decoding accuracy, given the responses from a brain region as input, and interpreting decoding with significantly-above-change accuracy as evidence that the brain region encodes information about the property that was successfully decoded [3]. Similarly, in the field of functional connectivity, the best estimate of the statistical dependence between the responses in two different regions is calculated, and significance is interpreted as evidence for the dependence between those regions’ responses [10, 11].

In this article, we introduce a new perspective. We suggest that nonlinear models should not be used to replace linear models – instead, information about the relative contributions of linear and nonlinear models should be preserved. Rather than selecting a single model and using its performance to determine the strength of the interaction between two regions, we propose to use a family of models and to treat the respective contributions of different models as a “fingerprint” that characterizes not just the strength, but also the type of interaction between regions. From a mathematical perspective, the proposed approach is rooted in seeing the problem of connectivity as function estimation, and it is inspired by the idea that a function can be expressed as a point in a Hilbert space, having as coordinates its projections on a (infinite) set of basis functions.

Previous research has introduced nonlinear approaches to the study of connectivity using Dynamic Causal Modeling (DCM, [12]), Granger causality [13], mutual information [14], and Multivariate Pattern Dependence (MVPD, [15]). However, by and large these methods have followed the traditional approach of building one model that performs as accurately as possible and interpreting the quality of fit, parameter values, or accuracy as evidence for the existence of interactions. The approach we propose in this article, instead, focuses on distinguishing between different kinds of interactions between regions, offering a new technique that can reveal differences even between region pairs whose overall correlations or statistical dependencies are comparable in strength.

## Methods

The study of univariate statistical dependence between pairs of brain regions offers an ideal test case for the use of computational fingerprints. The univariate nature of the problem prevents a combinatorial growth in the number of nonlinear terms, and the continuous (rather than discrete) nature of the outputs makes it possible to use a simple basis set such as Hermite polynomials ([16, 17]). We used Hermite polynomials in this work as they are defined on all ℝ and have a natural multivariate extension, but the same logic can be applied to other basis sets such as Fourier basis functions or Legendre polynomials. For additional convenience, in this study we divide the each Hermite polynomial of order *n* by 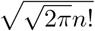 to render them an orthonormal basis (see [18], Appendix, The first five normalized Hermite polynomials, see also Figure 1, A).

**Figure 1:**
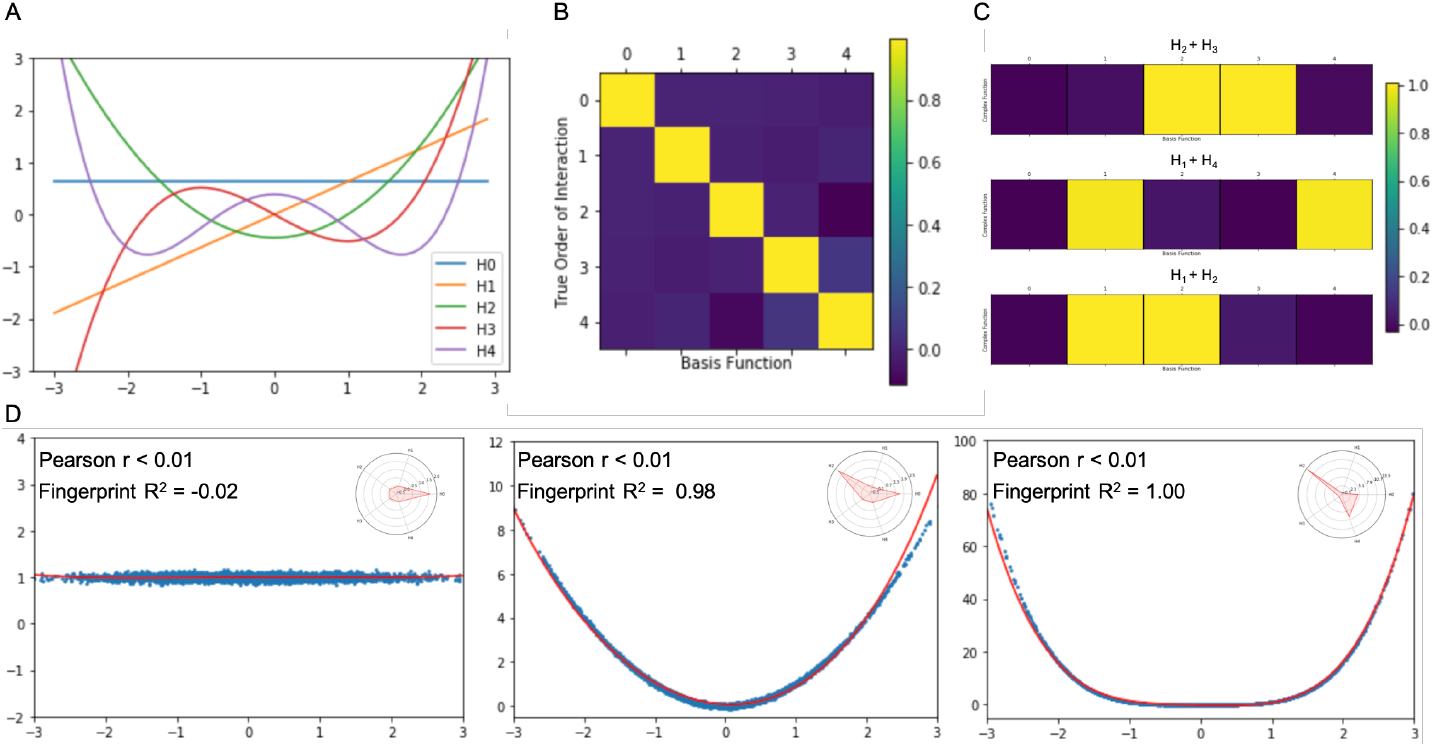
Validation with synthetic data. **A**. The first 5 Hermite polynomials. **B**. Estimated fingerprints for datasets generated following the first five Hermite polynomials. As expected, the estimates assign coefficients near 1 to the correct polynomial in the basis set, and coefficients near 0 to all others. **C**. When the data are generated using a linear combination of two basis functions, the resulting fingerprint reveals loadings near 1 for each contributing basis function, and loading near zero for all other basis functions. **D**. Nonlinear fingerprints can approximate u-shaped interactions that would be indistinguishable using standard correlation analysis.

## Theory

### Expressing functions with a truncated orthonormal Hermite basis

We will consider the average response in a predictor region at time *t* (which we will denote with *x*_*t*_), and the average response in a target region also at time *t* (which we will denote with *y*_*t*_). Modeling the dependence between *x*_*t*_ and *y*_*t*_ as *y*_*t*_ = *f* (*x*_*t*_) + *ϵ*_*t*_, we will aim to characterize the function *f* that minimizes the error 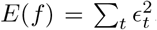. For convenience (and without loss of generality), we will normalize the inputs and outputs to have mean 0 and standard deviation 1. We will make the assumption that the function *f* is in the Hilbert space of functions from the interval ℝ to ℝ satisfying

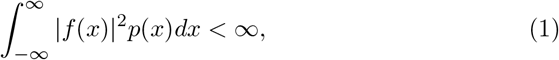

where

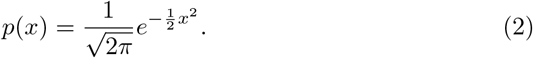

This is a large space of functions, and it should be sufficient to approximate well the relationship between the predictors and the targets of prediction.

The inner product between any two functions *g*_1_, *g*_2_ defined as

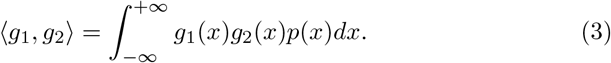

Hermite polynomials are a basis set of the Hilbert space. If we knew the function *f* that minimizes the error *E*(*f*), we could express it as an infinite set of coordinates *c*_*i*_ where

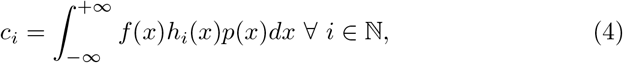

where *h*_*i*_ is the *i*th Hermite polynomial (normalized to have norm 1).

To make this strategy applicable in practice, we need to address two challenges. First, we can’t calculate an infinite number of coordinates - therefore, we will truncate Hermite polynomials to a specified order. The optimal order at which to truncate the polynomials can depend on the amount of data and the nature of the interactions between the regions studied. In the present article, the main focus is not to determine the optimal number of polynomials. Therefore, we will use polynomials up to the fifth order (future studies can use variance explained in independent data as a metric for the selection of the number of polynomials). Second, we do not know the function *f*, we only have a training dataset containing pairs of observations (*x*_1_, *y*_1_), …, (*x*_*T*_, *y*_*T*_). To address this second challenge, for each Hermite polynomial *h*_*i*_ we will estimate the corresponding coordinate as

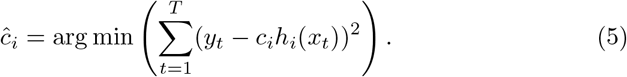

The coordinates *ĉ*_*i*_ approximately characterize the function *f* (up to the precision afforded by the truncation).

### Convergence of the coordinate estimates

If *ĉ*_*i*_ is a “good” estimate of *c*_*i*_, as the number of observations grows the estimate should converge to the true value *c*_*i*_. We demonstrate this property in the following Lemma.

#### Lemma 1.

*We will assume that the data generating process is approximately Normal, and since we normalize our input data to have mean µ* = 0 *and standard deviation σ* = 1 *we have approximately that x* ∼ *𝒩* (0, 1) *(see Appendix, Normality of the Data; Figure 7). Since the error function is convex, we can calculate ĉ*_*I*_ *by setting*

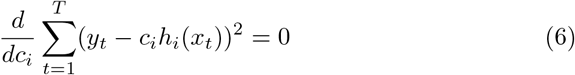

*which yields*

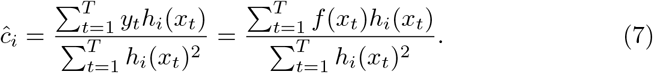

*As the number of observations increases, taking into account the fact that x* ∼ 𝒩(0, 1), *we have that*

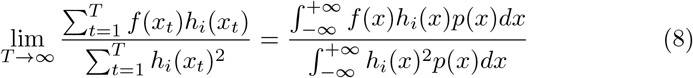

*and since we have used normalized Hermite polynomials that form an orthonormal basis*,

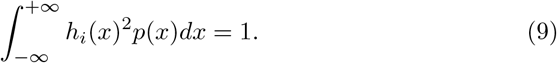

*In conclusion*,

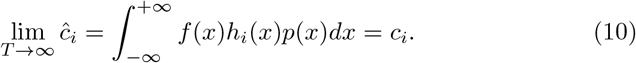

Note that based on this observation, in the presence of non-normally distributed data, it might be possible to estimate the probability density of the data *q*(*x*) and develop a basis set of polynomials that are orthonormal with respect to the inner product defined by

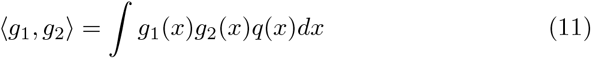

where the integral is computed over the domain of *q*.

## Application

### Participants and stimuli

All data used in this study were made publicly available as part of the *Study-Forrest* dataset [19]. These data consist of fMRI scans of 15 subjects (6 female, ages 21-39, mean = 29.4) as they watched the movie *Forrest Gump*. Data from one subject were removed from the analyses after failing the fMRIPrep preprocessing procedure [20] see also [21].

After providing consent, participants watched the movie in the scanner over the course of 8 functional runs (approximately 15 minutes each). Additionally, subjects performed a localizer task incorporating 24 grayscale images from each of the following six categories: faces, bodies (without heads), small objects, houses, outdoor scenes of nature and streets, and phase scrambled images; for more information about the localizer task, please see [22].

All scans were performed in a 3T Philips Achieva dStream MRI scanner with a 32-channel head coil. BOLD responses were recorded at 3 × 3 × 3mm resolution with T2*-weighted echo-planar (2 sec repetition time (TR)) imaging sequence. See [19] for more details on image acquisition.

### Data preprocessing

Data were preprocessed according to the fMRIPrep pipeline described in [20]. This procedure combines the following steps: T1-weighted (T1w) anatomical images were smoothed and skull-stripped using ANTs; brain images were segmented into white matter (WM), gray matter (GM), and cerebrospinal fluid (CSF) using FAST (FSL) [23]; MCFLIRT (FSL) [24] was used to correct functional scans for head-movement; functional scans were aligned with the corresponding anatomical image using boundary-based co-registration implemented in FLIRT (FSL).

After preprocessing, the data were additionally denoised using CompCor [25]. In this method, noise is removed from the functional data by regressing out the first five principal components extracted from the combined WM and CSF data.

### Data analysis

Let’s consider a fMRI dataset, and a seed region. In this study, we used the fusiform face area (FFA) as the seed region. We then used computational fingerprints to characterize the relationship between responses in FFA and responses in each other voxel *v* in gray matter. For each participant, and for each voxel in gray matter (normalized to MNI space), we applied the method described in the previous section using as *x*_*t*_ the average response in FFA at time *t*, and as *y*_*t*_ the response in the voxel *v* at time *t*. This procedure yielded a 5-dimensional vector of the estimated coordinates *ĉ*_1_, …, *ĉ*_5_ along the first 5 Hermite polynomials. Next, we used k-means clustering with the Akaike Information Criterion (AIC) to identify clusters of voxels using the 5-dimensional vectors from all participants. This approach identified the optimal number of clusters from the data by balancing complexity and quality of fit. Each cluster corresponded to a distinct kind of nonlinear interaction between brain regions. Finally, clustering was visualized by color coding each voxel in gray matter by the cluster to which it is assigned most often, with saturation increasing as a function of the proportion of participants for whom the voxel was assigned to the most frequent cluster.

To test our model’s ability to distinguish between voxels based on their nonlinear interactions with the FFA, we compared clustering solutions for the 5-dimensional fingerprints with the optimal clustering solution across the loadings for only the linear basis vector. In this way, we were able to determine the subsets of voxels with similar linear loadings that were differentiated by their nonlinear components. This analysis clusters voxels according to the *type* of interaction between the voxel’s activity and the activity in the seed region, thus highlighting brain areas with distinct functional relationships to the FFA. Finally, using Statistical Non-Parametric Mapping (SNPM) [26] we tested whether the magnitude of the nonlinear basis vector loadings for all voxels (using a cluster forming threshold of 0.0001) were significantly non-zero to determine where neural interactions with the FFA were significantly nonlinear in nature.

## Results

### Estimated coordinates match the ground truth in simulated data

In order to test the efficacy of our novel analysis to detect the functional relationship between two patterns of activity, we used simulated data to model a series of potential interactions between generated seed-target data sets. By manipulating the function used to create target data from a set of simulated seed data, we can test the ability of the fingerprint analysis to correctly model the selected relationship. Our results illustrate that through estimating loadings on the first 5 basis vectors of our selected functional space, we can accurately characterize the generative function of the target data for the functions tested (see Figure 1, more complex functions might require a higher number of polynomials).

#### Individual polynomials

Starting with a seed-sample of 10,000 normally distributed data points (mean = 0, SD = 1), we defined 5 sets of target data as *H*_1*−*5_(*seed*) where *H*_1*−*5_ represents each of the first 5 normalized Hermite Polynomials. Given that the interactions between the seed and target data were selected to be the 5 basis vectors by which we are measuring functional space, if our analysis correctly identifies the underlying computation that generates the target data, we would expect to see a loading of 1 on the relevant basis vector and loadings of 0 on all other basis vectors. For each set of target data, we found a 5-dimensional fingerprint with a loading of 1.00 for the basis vector governing the underlying relationship between the seed and target data, as well as loadings with an absolute value *<* 0.12 for all other basis vectors (see Figure 1, B).

#### Combinations of polynomials

To further probe the ability of a computational fingerprint to capture more complex relationships between seed and target data, we next generated target data using a linear combination of multiple Hermite polynomials. Assuming the underlying function that describes the interaction between the seed and target data is an unweighted combination of a subset of Hermite polynomials, the resulting fingerprint should consist of a vector of loadings with values of 1 for the given subset of polynomials and loadings of 0 for all other basis vectors. In our validation, we demonstrate in three cases (*H*_2_ + *H*_3_, *H*_1_ + *H*_4_, and *H*_1_ + *H*_2_) that these computational fingerprints capture the selective loadings on the relevant basis vectors (see Figure 1, C).

### Nonlinear fingerprints can capture U-shaped dependencies

One key benefit of characterizing functional interactions using nonlinear fingerprints is that they offer considerably more explanatory power than a standard Pearson correlation. One illustrative example in which our computational fingerprint analysis outperforms correlation tests arises in the instance when interactions are governed by a symmetrical underlying function. In the case of a U-shaped relationship, the magnitude of activity in a seed region, either negative or positive, results in a proportional positive response in the target region of interest. This type of relationship could be particularly useful to understand brain regions that might show greater sensitivity to the magnitude of the deviation from baseline of the responses in another brain region, as opposed to its direction (positive/negative).

Importantly, testing for these patterns of related activity using Pearson’s correlation will result in null findings for any significant interactions between the symmetrical data. Because Pearson’s R is a measure of the *linear* correlation of variables, it is not especially informative when seeking to explore *nonlinear* relationships between sets of data. In contrast to the inability of correlation coefficients to distinguish between null relationships and U-shaped dependencies, estimating a computational fingerprint to map interactions provides a much more informative model of any symmetrical dependencies (see Figure 1, D).

In our simulated experiments, we show that not only do fingerprint estimations tightly track the shape of nonlinear functions (e.g. *x*^2^*andx*^4^), but also, that these fingerprints are able to differentiate between dependencies that would otherwise be indistinguishable using measures of linear correlation(see Figure 1, D). To highlight the explanatory power of our fingerprint analysis, we generated 3 sample dependencies where Pearson R = 0: *y* = 1 + *ε, y* = *x*^2^ + *ε, andy* = *x*^4^ + *ε*, where *ε* represents a random amount of noise selected from a normal distribution (*mean* = 0, *SD* = 1) scaled by a factor of 0.05. Importantly, while all 3 of these interactions show *R <* 0.01, each relationship is described by a unique computational fingerprint (*y* = 1: [1.58, 0.01, 0.00, 0.00, 0.013]; *y* = *x*^2^: [1.61, −0.01, 2.26, 0.17, 0.1642599]; *y* = *x*^4^: [4.64, −0.04, 13.07, 0.61, 7.04]). In this way, these fingerprints allow for the identification and modelling of U-shaped dependencies that could otherwise be overlooked in a standard correlation analysis.

### Nonlinear fingerprints identify more clusters than linear fingerprints

Exploring neural data using computational fingerprints provides unique insight into the *types* of interactions between brain regions. Moreover, using additional, nonlinear basis vectors to estimate inter-regional dependencies allows for a heightened sensitivity to more complex interactions. Other approaches like mutual information can be used to capture nonlinear dependence between brain regions [14, 27]. However, computational fingerprints are unique in that they do not characterize the interaction between two brain regions using a single value that reflects the strength of the dependence; instead, computational fingerprints characterize the interaction between two regions with a multi-dimensional vector encoding the contributions of different Hermite basis functions. We used these multi-dimensional vectors to subdivide cortex into distinct clusters of voxels with different types of interactions with the FFA. To quantify the advantage of using multi-dimensional fingerprints, we compared the optimal clustering solutions for voxels across participants using 5 dimensional fingerprints (leveraging the first 5 normalized Hermite polynomials as basis vectors) and 1 dimensional fingerprints (using only the 1st order, linear polynomial). In both cases, we calculated the AIC for clustering solutions ranging from 1 to 10 clusters. In order to counterbalance the increased explained variance of more clusters with the potential to over fit the data with too many clusters, the optimal number of clusters is determined by locating the “elbow” of the plotted AIC values– where an increase in the number of clusters no longer corresponds with a substantial decrease in information lost. Importantly, for the linear fingerprint analysis, the optimal number of k-clusters was found at *k* = 2 while for the nonlinear fingerprint analysis, the best solution existed at *k* = 5 (see Figure 2, A and C). After determining the ideal k value for the linear and nonlinear analyses, we used k-means to group voxels by their linear fingerprints and nonlinear fingerprints across all subjects (see Figure 2, B and D). Our analysis of the linear fingerprints yielded 2 distinct clusters across the brain. The smaller (yellow in Figure 2, B) of the two clusters encompasses the dorsal and ventral temporal lobes bilaterally, as well as large sections of the bilateral visual cortex. The larger cluster (blue in Figure 2, B), comparatively, sprawls across the frontal and parietal lobes, as well as the lateral temporal lobes in both hemispheres. In contrast, the 5 clusters generated from the nonlinear fingerprints show a unique division of the cerebral cortex (Figure 2, D). The green cluster is concentrated around a series of face selective regions, including the FFA (our seed region), the superior temporal sulcus (STS), and the occipital face area (OFA). The purple cluster covers the majority of the visual cortex as well as the lateral and ventral temporal lobes. The red cluster is more distributed, encompassing bilateral sections of the lateral temporal lobes, early visual cortex, as well as sparse sections of the prefrontal cortex (PFC) and medial frontal lobe. The yellow cluster is located in 4 distinct areas associated with the default mode network (DMN): the PFC, the precuneus, the angular gyrus, and the lateral temporal cortex. Finally, the blue cluster spans large sections of the frontal and parietal lobes, as well as more sporadic areas in the anterior and lateral temporal lobes.

**Figure 2:**
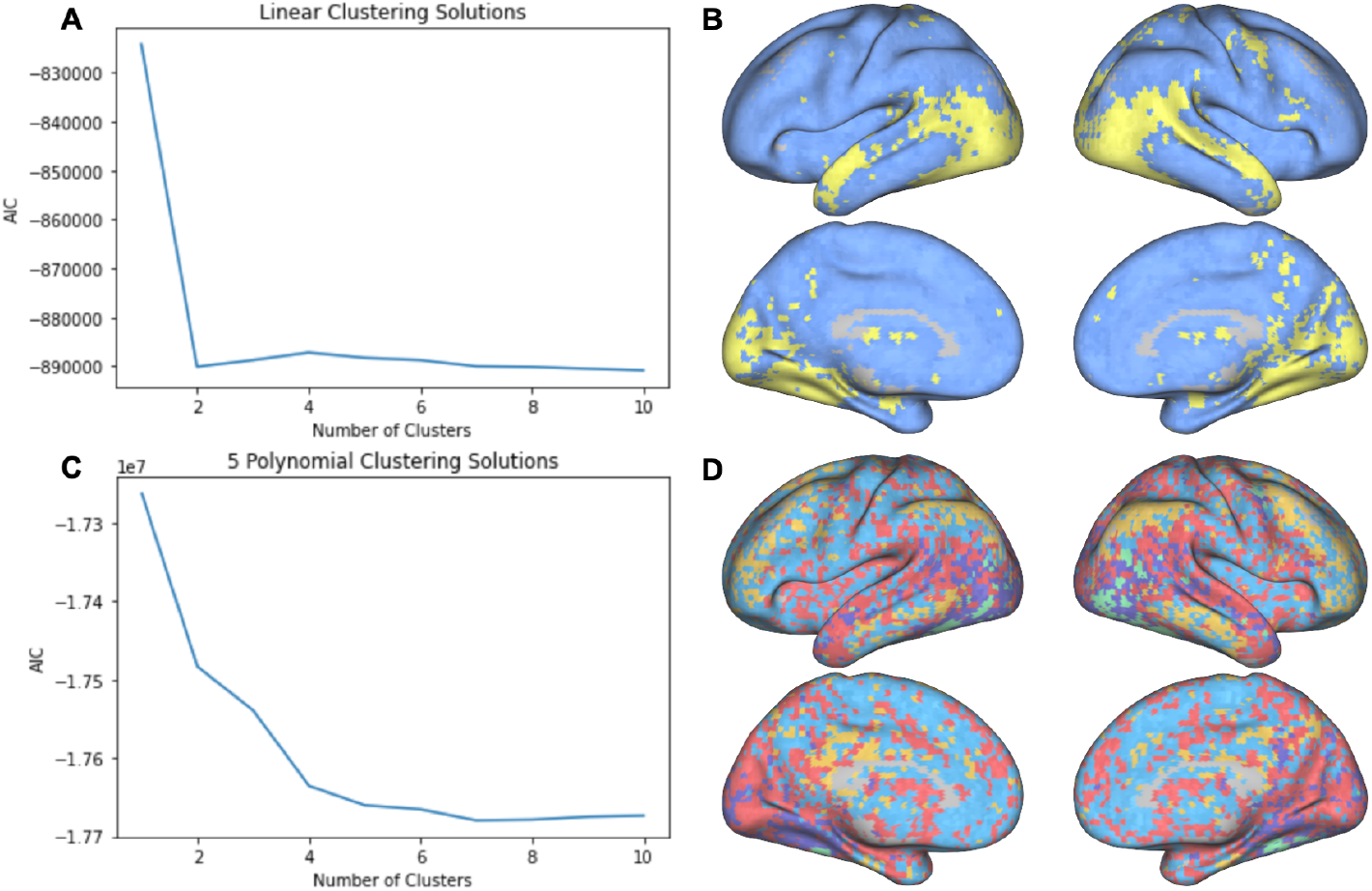
**A**. Plot of Akaike Information Criterion (AIC) values for K-means clustering solutions with different numbers of clusters using linear fingerprints (Hermite polynomials of degree 1). The elbow occurs at *k* = 2 clusters. **B**. Visualization of the cluster assignments of different voxels based on the linear fingerprints. For each voxel, we used the mode cluster assignment across subjects to determine that voxel’s final cluster value. Each cluster is represented by a different color and the intensity of the color represents the percentage of subjects sharing that voxel’s cluster assignment (lighter colors denote a higher percentage). **C**. Plot of AIC values for K-means clustering solutions with different numbers of clusters using nonlinear fingerprints (Hermite polynomials of degree 0 through 4). The elbow occurs at *k* = 5 clusters. **D**. Visualization of the cluster assignments of different voxels based on the nonlinear fingerprints (generated with the same approach described in B).

### Distinct clusters are associated with unique functional relations to the FFA

One of the central advantages of the fingerprint analysis is an enhanced interpretability of nonlinear dependencies. Because the nonlinear components of the estimated function are quantified as loadings on basis vectors, we can calculate the interaction between two regions as a weighted sum of the normalized Hermite polynomials. To this end, we can observe the unique computational relationship that defines a given cluster, by taking the loadings from the k-means defined cluster center (see Figure 3). After segmenting the grey matter voxels using k-means clustering, we identified 5 distinct clusters with centers at: [4.34*e*−09, 4.78*e*−01,−3.34*e*−02, 2.23*e*−02,−2.87*e*−02], [4.19*e*− 09, 2.61*e*−01,−9.47*e*−03, 2.43*e*−02,−1.30*e*−02], [4.28*e*−09, 1.35*e*−01,−6.60*e*− 04, 1.51*e*−02,−4.85*e*−03], [4.32*e*−09,−8.28*e*−02, 8.20*e*−03,−7.01*e*−05, 6.32*e*− 03], [4.36*e* − 09, 3.40*e* − 02, 6.56*e* − 03, 5.40*e* − 03, 1.46*e* − 03]. For each cluster, we calculated the defining function using the respective central fingerprint as the loadings on the basis vectors.

**Figure 3:**
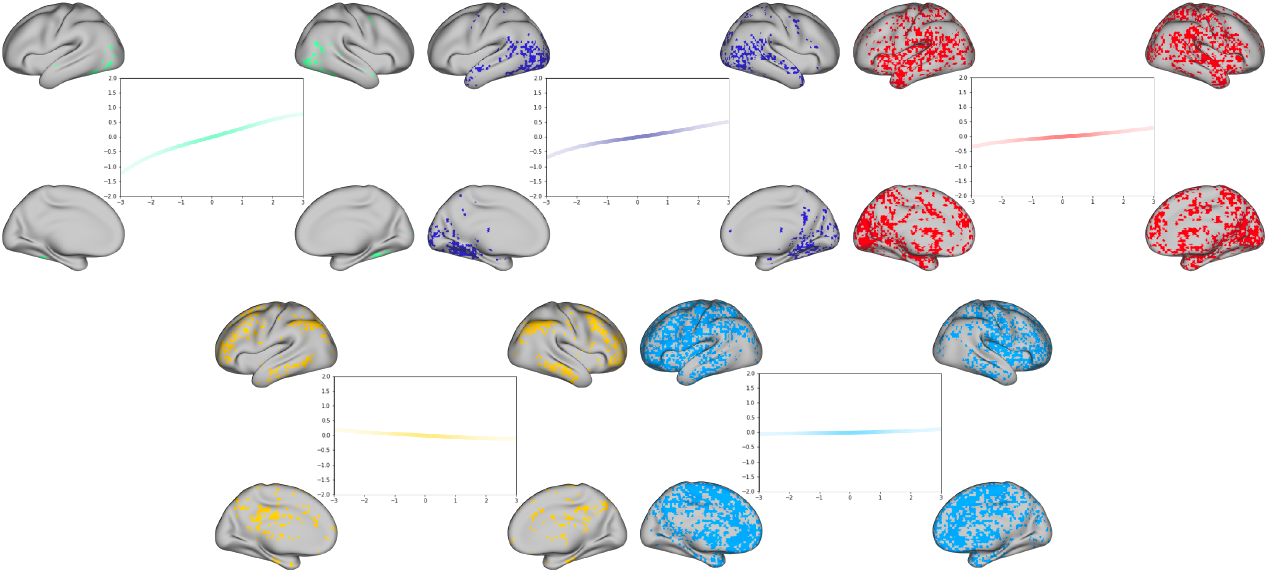
Clusters of voxels with different associated fingerprints. The strongest positive relationship is observed in ventral and lateral posterior temporal regions (green cluster). Note a negative relationship with a cluster of regions in the vicinity of the default-mode-network (in yellow cluster).

### Clustering solutions show anatomical consistency with the increase in the number of clusters

In order to test the impact of selecting different k values on the spatial layout of the resulting clusters, we re-ran the k-means clustering using k values ranging from 2-5 (see Figure 4). Importantly, we found that the clustering solutions were consistent in their groupings of key areas across the inferior and superior temporal lobe, the visual cortex, as well as the PFC. This consistent anatomical grouping across clustering solutions suggests that our findings are robust across clustering solutions and not dependent on the selection of a distinct number of clusters.

**Figure 4:**
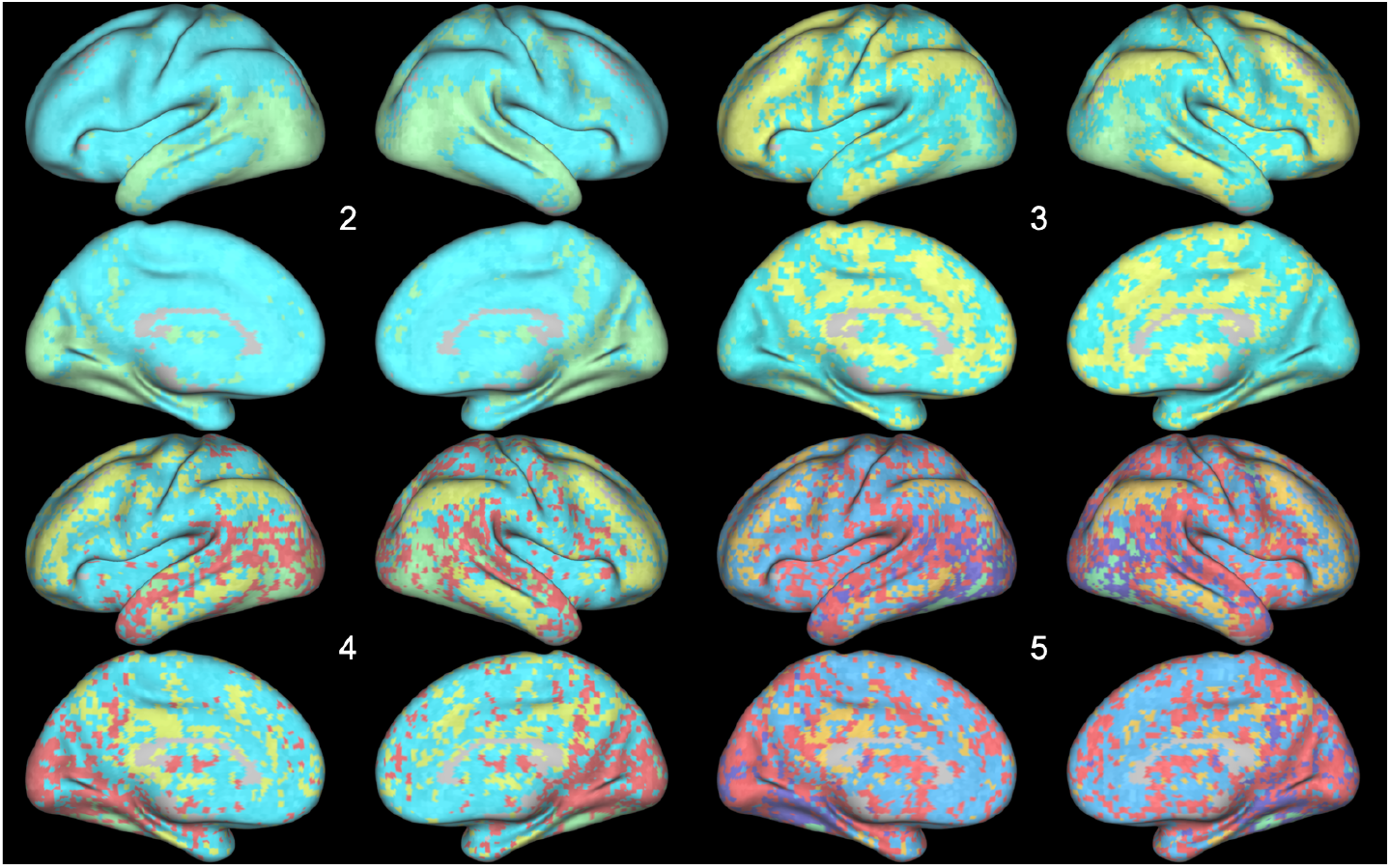
K-means clustering of voxels based on 5-dimensional fingerprints, for values of K ranging from 2 to 5 clusters. As the number of clusters increases, the spatial layout of the new clusters provide a more detailed parcellation of the cortex, but highlight a strikingly similar set of regions.

### Coordinates along dimensions of the Hilbert space reveal the distribution of nonlinearities across cortex

In order to observe the distribution of nonlinear interactions across the cortex, we next plotted the loadings for each of the individual basis vectors for each voxel (see Figure 5, additionally see the appendix Figure 8) for the loadings on the 1st basis vector). Interestingly, while the loadings for the linear basis vectors were highest in the regions surrounding the face selective cortical regions and dorsal temporal lobe (Figure 5, 1st Order), nonlinear loadings, particularly in in the 2nd and 4th order basis vectors, were highest in frontal and medial regions, with negative loadings in the face selective cortex and lateral temporal regions. It is worth noting, that the magnitude of the basis vector loading reflects the contribution of that basis vector to the overall relationship between the given voxel and the FFA. For this reason, larger loadings on nonlinear polynomials, reveal stronger nonlinear components within the given interaction.

**Figure 5:**
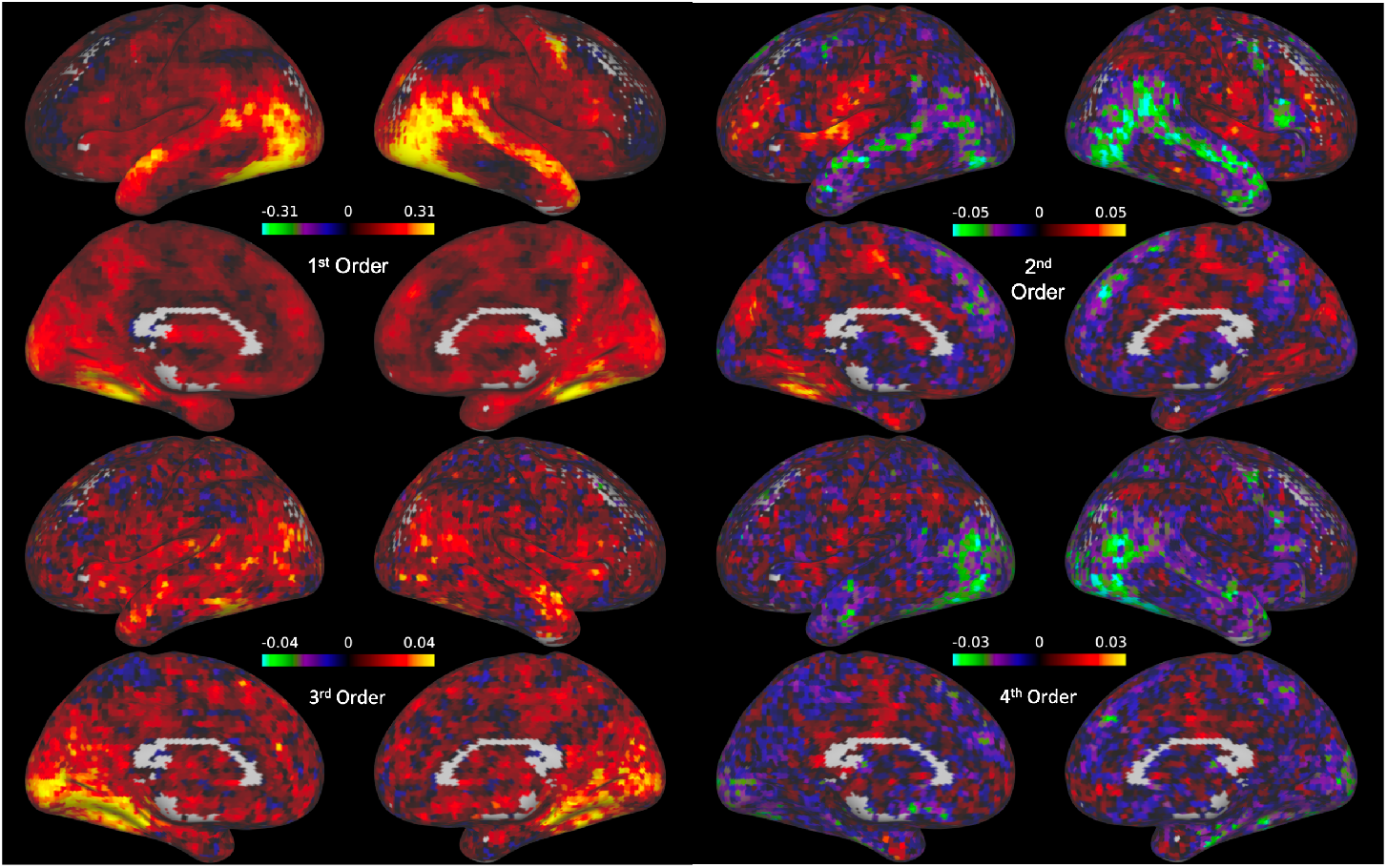
Whole-brain maps of the coefficient estimates for the Hermite polynomials in the basis set (excluding the constant 0th-order polynomial, see Figure 8). Different polynomials in the basis set are associated with unique cortical maps of coefficients. The maps of the 2nd order and 4th order basis vector loadings demonstrate an increased magnitude of the loadings for their respective polynomials in the dorsal temporal lobe and medial frontal lobe, as well as the face selective visual cortex.

To further test the significance of these nonlinear interactions across the cortex, we performed across-subject one-tailed T-tests using SnPM to highlight any clusters of voxels with significantly non-zero loadings on each of the nonlinear basis vectors (i.e. the 2nd, 3rd, and 4th order basis vectors, Figure 6). These tests revealed one cluster of voxels in area V5 with significantly negative loadings on the 2nd order basis vector (cluster threshold *p* = .0001, *p*(*FWEcorrected*) *<* .025). Additionally, we found 24 clusters in the early visual cortex and medial temporal lobe with positive loadings on the 3rd order basis vector (cluster threshold *p* = .0001, all p values (FWE corrected) *<* .025). We found no significant clusters with non-zero loadings on the 4th order basis vector.

**Figure 6:**
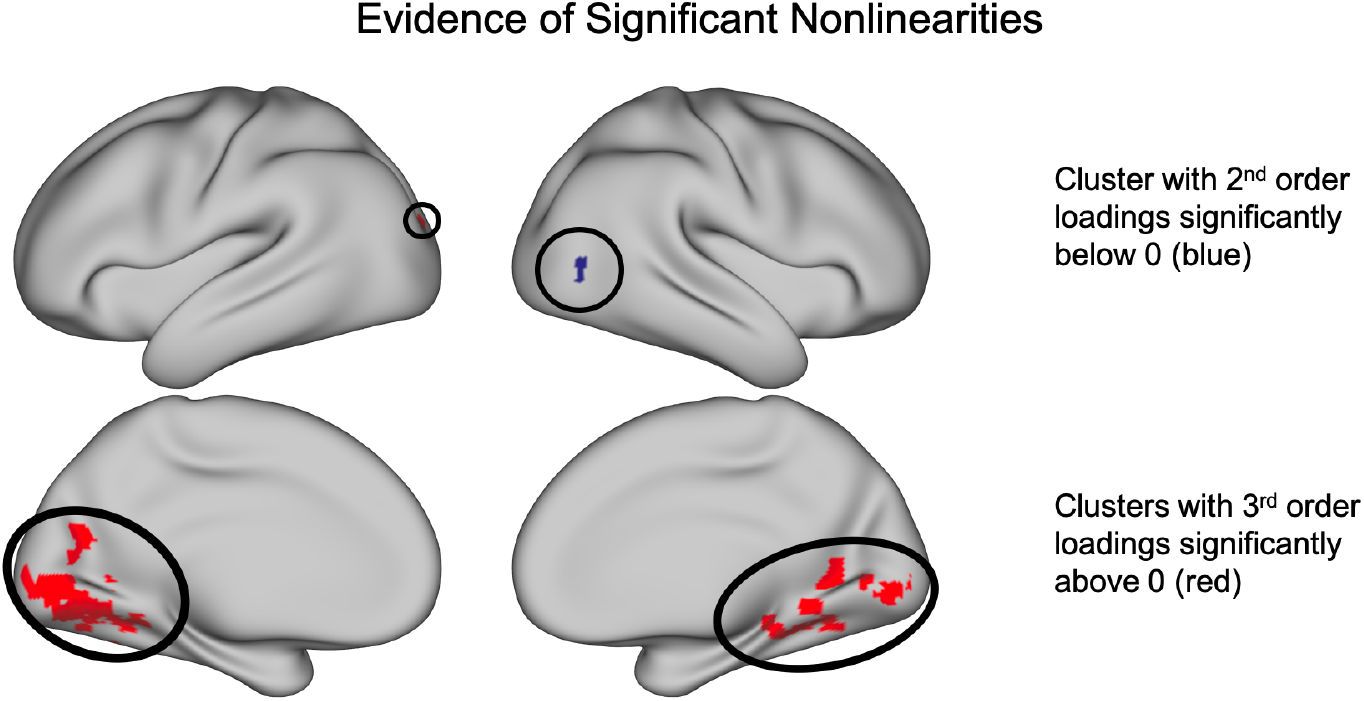
Significant clusters of non-zero loadings (SnPM *t*(13) ≥ 5.59, *p <* 0.05 FWE corrected). T-tests revealed a significant cluster of voxels with negative loadings on the 2nd order basis vector as well as a collection of clusters in the medial occipital and temporal lobes with positive loadings on the 3rd order basis vector.

## Discussion

### Computational fingerprints provide novel insight into neural interactions

The major contribution of this research is a novel method to investigate nonlinear interactions between brain regions: “computational fingerprints”. Understanding the complexity of inter-regional connectivity is an important objective within the landscape of current neuroscience research [15, 28, 12]. Importantly, computational fingerprints allow for a balance between the interpretability of linear models of connectivity [29] and the increased explanatory power and biological plausibility of nonlinear models [15]. Moreover, our method offers the ability to characterize *how* the responses in different cortical regions are related, and not just *whether* they are related.

Here, we demonstrated the capacity of the fingerprint model to capture various types of nonlinearities using synthetic data. Importantly, computational fingerprints were able to identify and distinguish between complex nonlinear relationships between data including symmetric (u-shaped) functions, which are particularly difficult to identify using only linear methods. This success on artificial data is an important indicator of the model’s potential to identify the existing nonlinear interactions in real functional data.

Common methods used to study cortical networks rely on linear tools that are incapable of capturing the wide array of potential interactions between brain regions. Computational fingerprints can capture the same linear interactions as methods like functional connectivity [30, 5]. However, using the loadings from the nonlinear basis vectors, it is additionally possible to extract the *nonlinear* components of a given relationship.

Some methods capable of capturing nonlinear interactions do exist [14, 27]; however, they often focus on quantifying the *strength* of statistical dependence, rather than studying the *type* of dependence. Importantly, the responses in different target brain regions could be predicted equally well by a given predictor region, yet at the same time, be related to the predictor region via different functions. Computational fingerprints are able to quantify the contribution of each nonlinear component of the overall functional relationship, allowing (unlike mutual information) for the explicit definition of the function as a whole (see Figure 3).

In some respects, computational fingerprints are related to estimating a polynomial fit of the interaction between two brain regions and using the vectors of coefficient estimates to characterize the interaction (to our knowledge, this approach has not been proposed in the previous literature). However, computational fingerprints offer a key advantage over using the coefficients from a polynomial fit. This is because in a polynomial fit, including higher-order nonlinear terms would also lead to changes in the coefficient estimates for the lower-order polynomials, making it difficult to compare the results across different studies that used models of different order. By contrast, due to the orthogonality of Hermite polynomials, computational fingerprints are such that the estimates of the coefficients of lower-order basis functions do not change when the model is expanded with the addition of higher-order basis functions.

Moreover, computational fingerprints can also be scaled up to take multidimensional patterns of activity as input, much like existing methods such as MVPD [11] and Informational Connectivity (IC) [31], through the use of multivariate Hermite polynomials. This advantage allows for an examination of the rich high dimensional nature of neural data and provides the opportunity to incorporate a wider array of neural response patterns into the study of nonlinear interactions between brain regions.

### Evidence of nonlinear interactions between brain regions

In this work, we also demonstrate the existence of distinct nonlinear interactions that characterize the relationships between the FFA and other regions across the brain. After first defining a fingerprint to describe the relationship of the average FFA activity with the activity in each other grey matter voxel, we were able to cluster the fingerprints across all voxels to identify 5 distinct sets of voxels with different functional interactions (see Figures 2, D and 3. Importantly, these clusters differ from those identified with a linear model of connectivity (see Figure 2, B) highlighting the increased ability to discriminate between voxels based on their relationship to the seed region.

Upon mapping out the functions associated with each cluster (Figure 3), we were able to determine the functional forms of the relationships between activity in the FFA and activity across the rest of the brain. A key feature of these relationships is their nonlinear components. While the loadings on the linear basis vectors are, indeed, still the predominant component in the defined clusters, the loadings on the nonlinear component are also important for differentiating the patterns of connectivity across voxels. This is apparent in the increase in the optimal number of clusters calculated for the 5 dimensional fingerprint (5 clusters) as compared to the clusters for the linear loadings only (2 clusters). Noticeably, voxels surrounding face selective regions and those in the anterior temporal lobe were grouped into a single cluster when only including the linear component; in contrast, when clustering across all 5 basis vectors, this same group of voxels was, instead, classified into separate, functionally distinct clusters. Moreover, we report additional evidence for the importance of discovered nonlinearites in the significant clusters of positive loadings on the 3rd order basis function and negative loadings on the 2nd order basis function (Figure 6). These clusters demonstrate specific areas in both early visual cortex as well as downstream visual processing regions with which the FFA has significant nonlinear interactions. In a follow-up analyses (see Appendix, Regressed Non-linear Loadings) we regressed out the linear basis vector loadings from those of the nonlinear basis vectors in order to discover any nonlinearities whose spatial distribution across cortical voxels is decoupled from that of linear interactions. With this analysis, we found two additional clusters that were trending towards significance in the posterior superior temporal sulcus (pSTS) and the anterior temporal lobe (Appendix, Figure 9), two regions previously implicated in face perception [32, 33].

These clusters suggest the possible presence of very localized, anatomically specific nonlinear interactions between FFA and other regions in the face network - but additional studies will be needed to evaluate the robustness of this finding.

### Potential pitfalls in the search for nonlinearities

Although, in this work, we present evidence of significant nonlinear interactions between brain regions, it is important to note that the linear components of these interactions were still an order of magnitude larger than the nonlinear loadings. Moreover, while we did find significant clusters of non-zero nonlinearities, these regions were particularly small and highly localized. These results are counterintuitive to the widespread nonlinearities that seem to be necessary given the existing computational and biological evidence [34, 35, 6]. For example, a recent study found that approximating the input/output relationship of a single pyramidal neuron requires using a deep recurrent neural network with 5 to 8 layers [7]. This apparent contradiction lends an important question for research moving forward: why do nonlinear interactions not account for a greater proportion of the variance in the interactions between brain regions?

The first potential explanation of the dearth of significant nonlinear interactions is that our model is incapable of identifying the full extent of the existing nonlinearities in the relationships between BOLD activity in different regions. However, our simulation data show that the method is able to accurately capture nonlinearities, at least within the set of cases we tested. In addition, it is important to note that we *do* report evidence of significant nonlinearities between the FFA and early visual cortex, V5, and medial temporal areas (see Results, Coordinates along dimensions of the Hilbert space reveal the distribution of nonlinearities across cortex). This suggests that our model is *capable* of capturing existing nonlinear interactions, but that the magnitude of these interactions in the fMRI data we analyzed is small. Additionally, our current model has outperformed previous models attempting to capture the nonlinear interactions with the FFA within this same dataset [36]. In sum, the small effect size of the nonlinearities appears to be a feature of the data, and not a consequence of the chosen model.

Another possibility that could explain the difficulty in discovering nonlinear interactions is that fMRI data, in particular, is not suitable to measure them. Previous work in this area has demonstrated that contributions of nonlinear models of functional connectivity in resting state fMRI have been relatively minor when compared to their linear counterparts [37]. This result could be, in part, due to a number of underlying limitations of fMRI data, including spatial resolution, temporal resolution, and hemodynamic smoothing.

When considering the mechanics of connectivity at a neural level, it is entirely possible that the nonlinear dynamics that we might expect to see are evident at a spatial scale that is too fine-grained for the millimeter resolution of an MRI scanner. At the cellular level, neurons are able to nonlinearly integrate signals from multiple synapses [6, 7]. In contrast, BOLD signal at the voxel level reflects the activity of hundreds of thousands of neurons. In this way, spatial smoothing might “blur” the complex interactions occurring at the neuronal level and thus, minimize the underlying nonlinear interactions. In a similar fashion to the spatial smoothing, fMRI data also suffers from temporal smoothing: our data might not be sampled at a high enough rate to map any nonlinear interactions that occur on the order of milliseconds. However, the averaging of a large number of nonlinear functions does not generally produce a linear function. If the loss of nonlinear information in fMRI signal is due to averaging over space and time, this would suggest that the amount and type of nonlinear interactions between neurons in different brain regions are such that their average is approximately linear.

Finally, another complicating factor of fMRI data is smoothing as the result of the hemodynamic function. Neural activity is nonlinearly related to the BOLD signal [35]. This proxy measurement of the underlying activity of cortical neurons can serve, in this case, as an additional smoothing function that might decrease the detectability of any nonlinear neural interactions.

Despite the possibility that the BOLD signal may not be the optimal tool for the study of the fine-grained nonlinear interactions between brain areas – it is also distinctly possible that it not the *quality*, but the *quantity* of data that is at issue. Hlinka et al. [37] only reported finding “subtle” nonlinear effects by, “testing across many pairs or even across many sessions.” In this study, we also found “subtle” yet significant nonlinear interactions by testing across a series of subjects. One potential avenue for future research might be to examine datasets comprising extensive scans of small numbers of participants [38]. Using many runs of single subject data and avoiding the averaging across subjects could increase the chances of identifying more subtle effects.

Although it is not currently clear why the scale of nonlinear interactions between brain regions is comparatively much smaller than the scale of linear interactions, we believe that solving this problem is *essential* for progress in the field of computational neuroscience. It is clear that nonlinear computations are crucial for models that approach human behavior on cognitive tasks [39], and for models that capture the behavior of biological neurons [7]. Applying methods such as computational fingerprints to very large fMRI datasets, or to datasets acquired with more direct measures of neural activity (i.e. neuropixels, [40]) is likely to shed new light onto the neural bases of cognitive computations.

## Code and Data Availability

The data used in this paper are from the public *StudyForrest* dataset and can be accessed at https://www.studyforrest.org/. The code to estimate the computational fingerprints can be found at https://github.com/cposkanzer/Computational-Fingerprints.

## Appendix

### The first five normalized Hermite polynomials

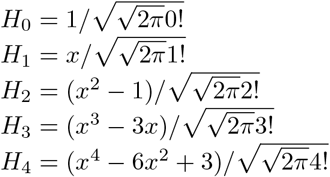

### Normality of the Data

See Figure 7

**Figure 7:**
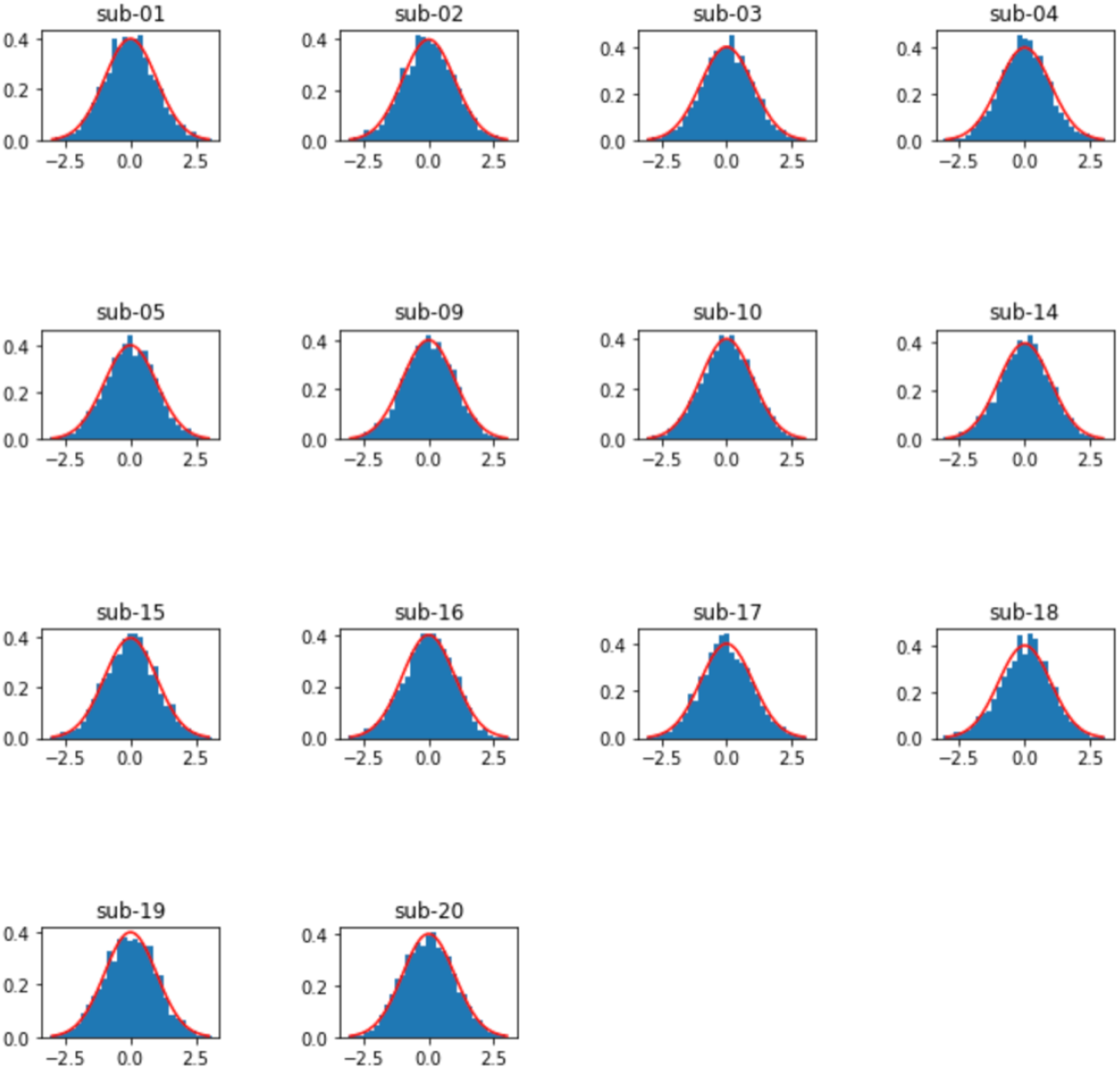
Z-scored data from each subject are plotted as probability density histograms (blue) with the the probability density function of the normal distribution (red) superimposed.

### Regressed Nonlinear Loadings

In addition to our analysis of the significance of the loadings on the nonlinear basis functions, we further probed the significance of the nonlinear loadings after regressing out the loadings from the linear basis function. We performed this analysis in order to to observe any nonlinearities that were uncorrelated with the linear interactions. We found 2 clusters of loadings on the 2nd order basis function that trended towards being significantly below zero: one in the pSTS (p = 0.04, FWE corrected one-tailed t-test), and one in the anterior temporal lobe (p = 0.04, FWE corrected one-tailed t-test) (Figure 9).

**Figure 8:**
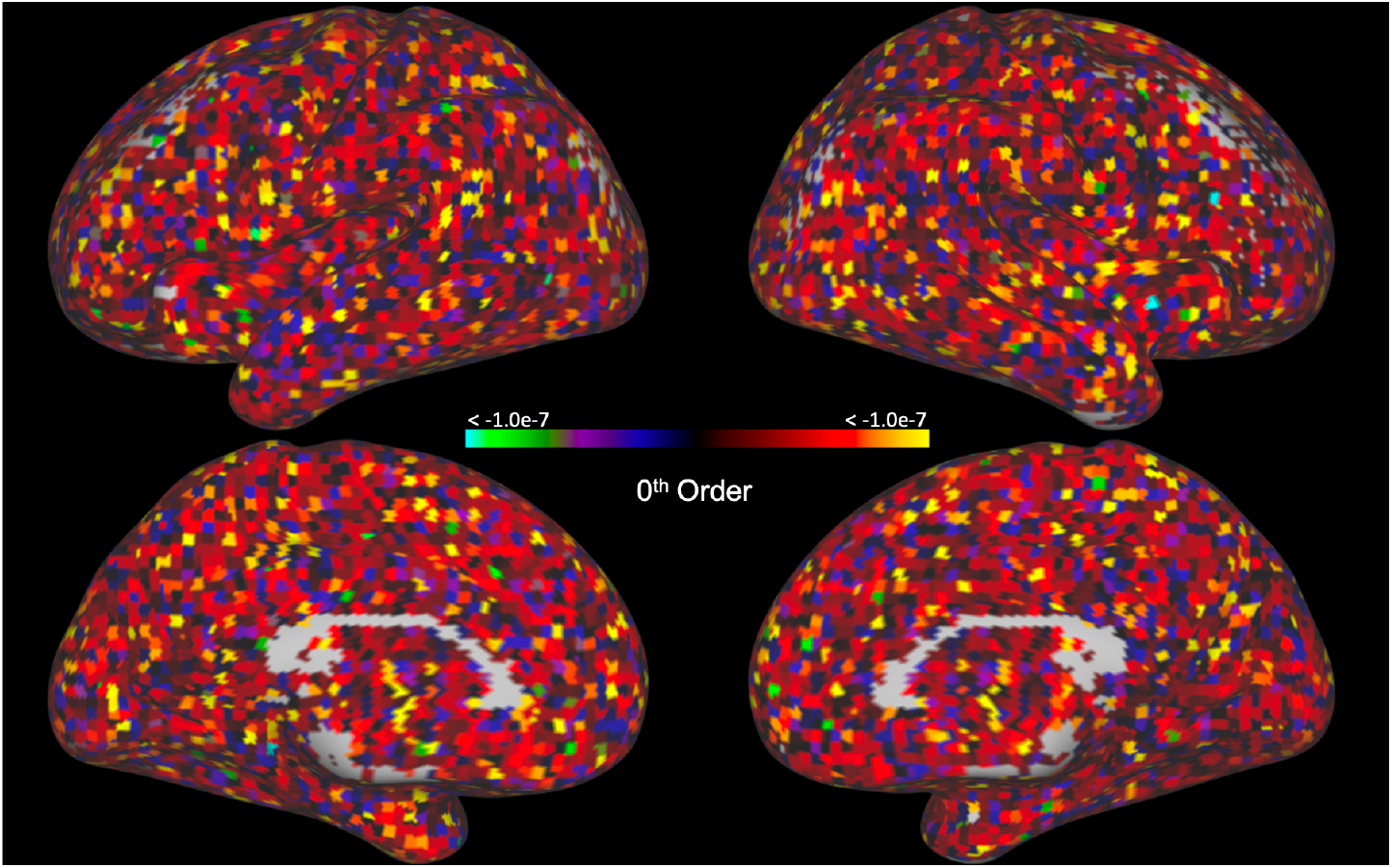
Distribution of loadings for he 0th order Basis function.

**Figure 9:**
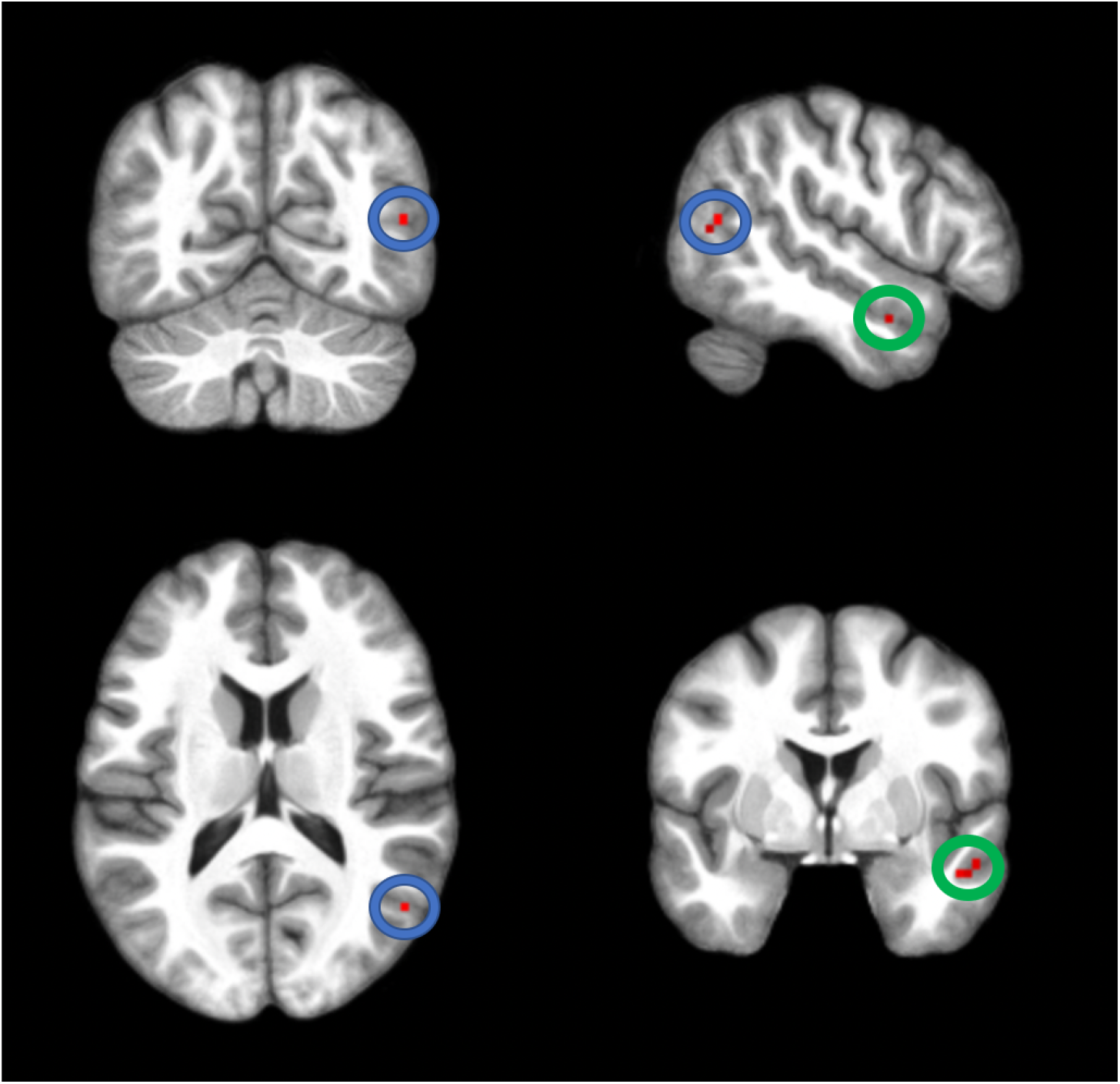
Clusters of below-zero loadings on the regressed 2nd order basis function.

